# Geometrical Study of Virus RNA Sequences

**DOI:** 10.1101/2021.09.06.459135

**Authors:** Alex Belinsky, Guennadi Kouzaev

## Abstract

In this contribution, some applications of the earlier developed fast algorithm of calculating coordinates of single nucleotides and RNA fragments are considered to create multi-scale geometrical models of RNAs and their mutations. The algorithm allows to plot single nucleotides and RNA’s fragments on one figure and to track the RNA mutations of any level visually and numerically using interpolation formulas and point-to-point estimates of coordinates of ATG starting triplets and single nucleotides. The performed study of many samples of SARS CoV-2 viruses shows perturbations of ATG starting triplet coordinates in the vicinity of *orf1ab* gene end only.

## 1. Introduction

The natural disaster is still ruling humankind over the globe. Although there is a great success in vaccines’ developments, nobody can guarantee new lines of known viruses or bursts of new infections, including bacterial or virological ones. The study of genetic information and how this information is transferred to the higher protein levels is a vital problem requiring joint efforts of biologists, medical doctors, gene researchers, and specialists from computer science, molecular and mathematical physics, etc. This short communication is with one of the methods of plotting RNA fragments along a genomic sequence for visual, quantitative, and statistical analyses of viruses threatening the world, and it that can be applied to any such species.

Many previous works in this matter relate to the beginning of this century [1]. They are with the imaging a single nucleotide in 4-D space or 21-D space along with a DNA/RNA or protein. Another approach is to present a sequence as a message with nucleotide symbols with the following quantitative modeling and apply one or several rather sophisticated signal processing techniques taken to dig out the properties of DNAs or RNAs [2].

## 2. Method

The proposed contribution here does not accentuate calculation technology. Still, unknown earlier features of the virus RNAs were found using a new technique based on a data mining algorithm combined with a signal processing method. A detailed calculation methodology can be found in [3], which is not considered in detail.

In short, the algorithm of searching fragments of RNAs is to represent a fragment and whole sequence by strings of binary numbers. The fragment moves along this sequence, and the Hamming distance value [4] is calculated between each binary representing these strings. If the compared characters are the same, then the result of Hamming distance calculation is zero. Otherwise, the product is expressed as a value between 0 and 1 and can be processed by a pertinent numerical algorithm.

Further, the code calculates the sub-sequences with the needed number of neighboring zeroes and their coordinates in the studied RNA sequence. The coordinates of the found fragments and single symbols can be placed together for multi-scale imaging on one plot for a more effective comparative analysis of RNAs and their mutations.

With these data, more operations are performed, and, for instance, the statistical properties of the codon length distributions are calculated. Among them are the medium values of codon lengths, r.m.s of these values, etc. As a new property of RNAs, in [3], the fractality of the codon length distributions and their type of fractality was found. The calculated fractal dimension value was connected to the virus family, and this dimension was relatively stable for the members of a virus branch.

The results discussed here are with more advanced imaging of SARS-CoV-2 and their comparative analyses following the initial research [3].

## 3. Results and Discussion

A virus sequence consists of a non-coding set of nucleotides at the beginning of a complete genome and several hundred codons. Codons start with triplets AU(T)G, where A stands for adenine, U for uracil, and G for guanine. Due to technical reasons, the uracil symbol U is substituted by T, which stands for thymine in the published sequence data. This substitution plays no role in symbolic and quantitative analyses of RNAs.

According to contemporary knowledge, mutations of viruses are with variations of RNAs due to environmental factors. They cause the substitution of single nucleotides in codons, vary their length, and change the codon’s number in RNAs. The role of a non-coding set is not clear at this moment, but it is a subject of mutations as well.

All mutations can lead to severe consequences of the virus properties because of re-building the capsid’s protein designs and replication rate of viruses. Anyway, the number of ATG triplets in one family of viruses is relatively stable, and it can be considered the RNA’s scheme or skeleton [3]. A study of re-building of these schemes is significant to search for the dramatic mutations of viruses. The proposed in [3] algorithm allows binding the ATG distributions together with the contents of each codon by the symbols of four mentioned nucleotides.

### 3.1. Multi-scale Mapping

Consider this mapping on an example of complete RNA of a sample of SARS CoV-2 virus from Wuhan (China) registered as MN988668.1 in GenBank [5] on Feb. 11, 2020. It counts 29881 nucleotides and 725 ATG triplets. Graphical representation of this RNA will help understanding its structure and possible mutations.

In Fig. 1, a map of this sequence is shown. On the abscissa, the coordinates of each nucleotide along the sequence are given. Each type of nucleotide (See Fig. 1’s legend) has its number placed on the ordinate axis. Then, this map allows obtaining the exact position of each nucleotide. By vertical lines, the coordinates of A symbols in ATG triplets are shown. Then, the space between these symbols is filled by TG +codon nucleotides.

**Fig. 1.**
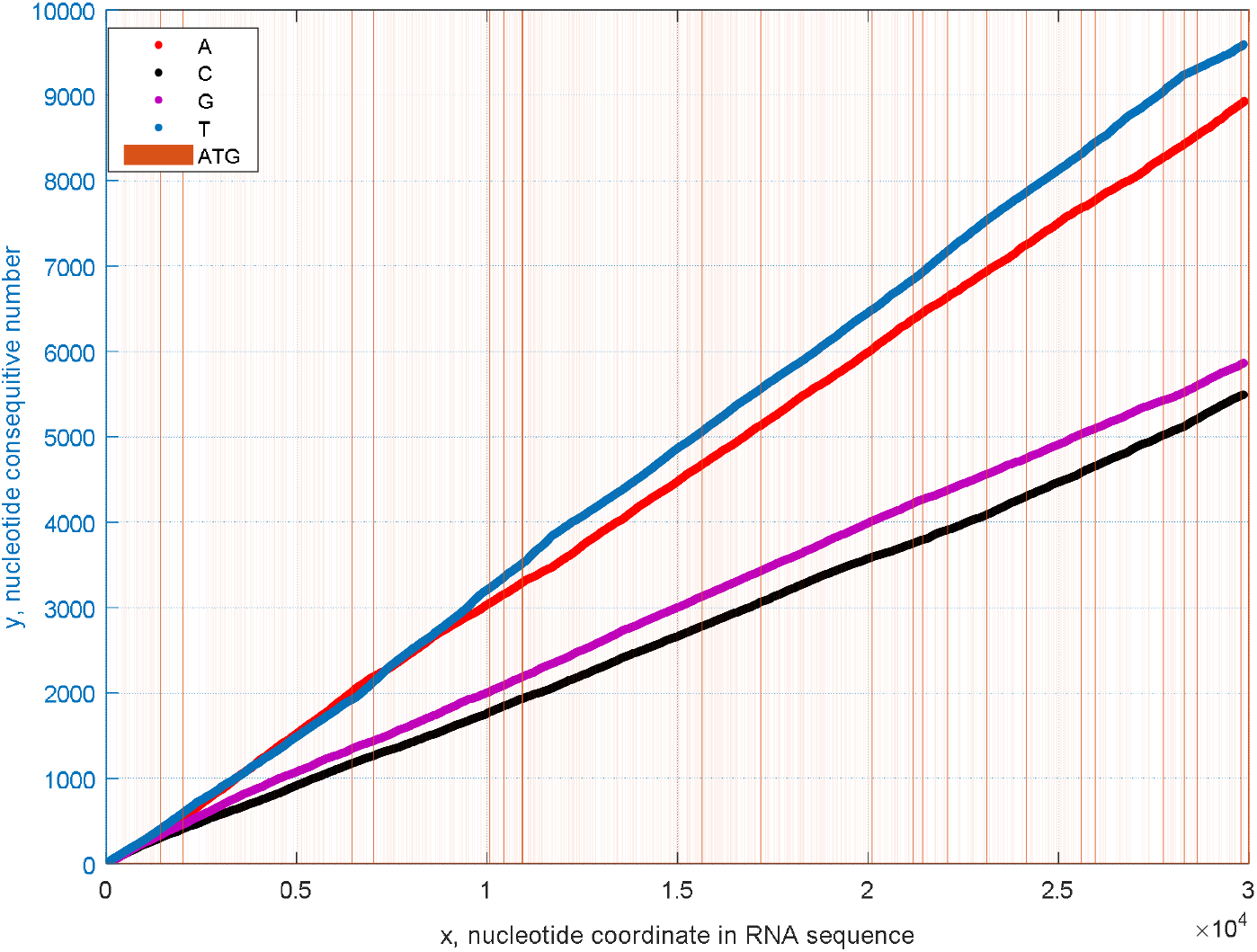
Map of the complete RNA sequence of MN988668.1 (GenBank) SARS CoV-2 virus sample.

Contemporary plotting tools allow interactive or programmed magnification of plots to watch for details. For our simulation, the Matlab Mathworks ® is used. The detailed map for the first 300 nucleotides is given in Fig. 2. It is seen that the first 105 nucleotides belong to a non-coding set. The first codon is then between the A symbols numbered 106 and 265, correspondingly. A colored point shows each nucleotide.

**Fig. 2.**
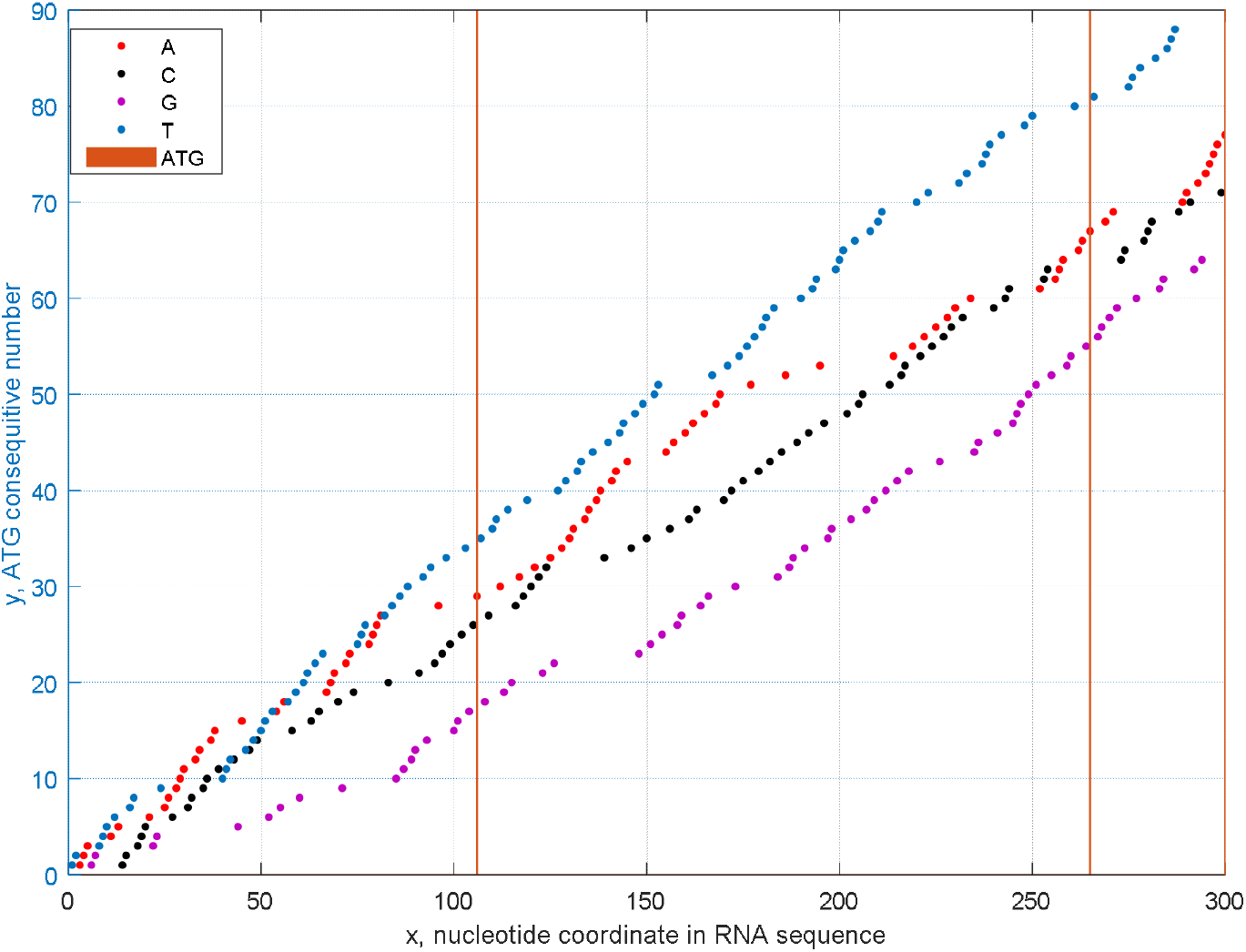
Magnified map of the complete RNA sequence of MN988668.1 (GenBank) SARS CoV-2 virus sample for the first 300 nucleotides.

### 3.2. RNA Interpolation Models

It is seen that the nucleotides are placed somewhat randomly. Their positions can be interpolated with some error by curves using the Matlab Curve Fitting Toolbox. The most straightforward interpolating formulas are shown below, and they allow calculating positions of symbols in 95% confidence bounds using linear polynomials:

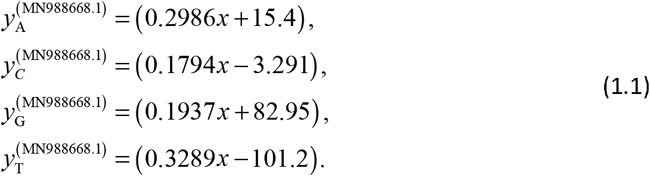

Similarly to the A, C, G, and T curves, the ATG dependence can be built, and a formula of the similar accuracy fits it:

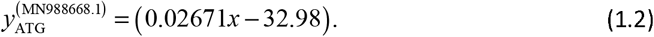

The formulas obtained here can be used for accelerated RNA modeling and estimates after careful study of their accuracy, considering Fig. 2. A graphical example of this interpolation is shown in Fig. 3. The mentioned tool allows interpolating data with better accuracy using polynomials of higher-order, spline technique, and other models.

**Fig. 3.**
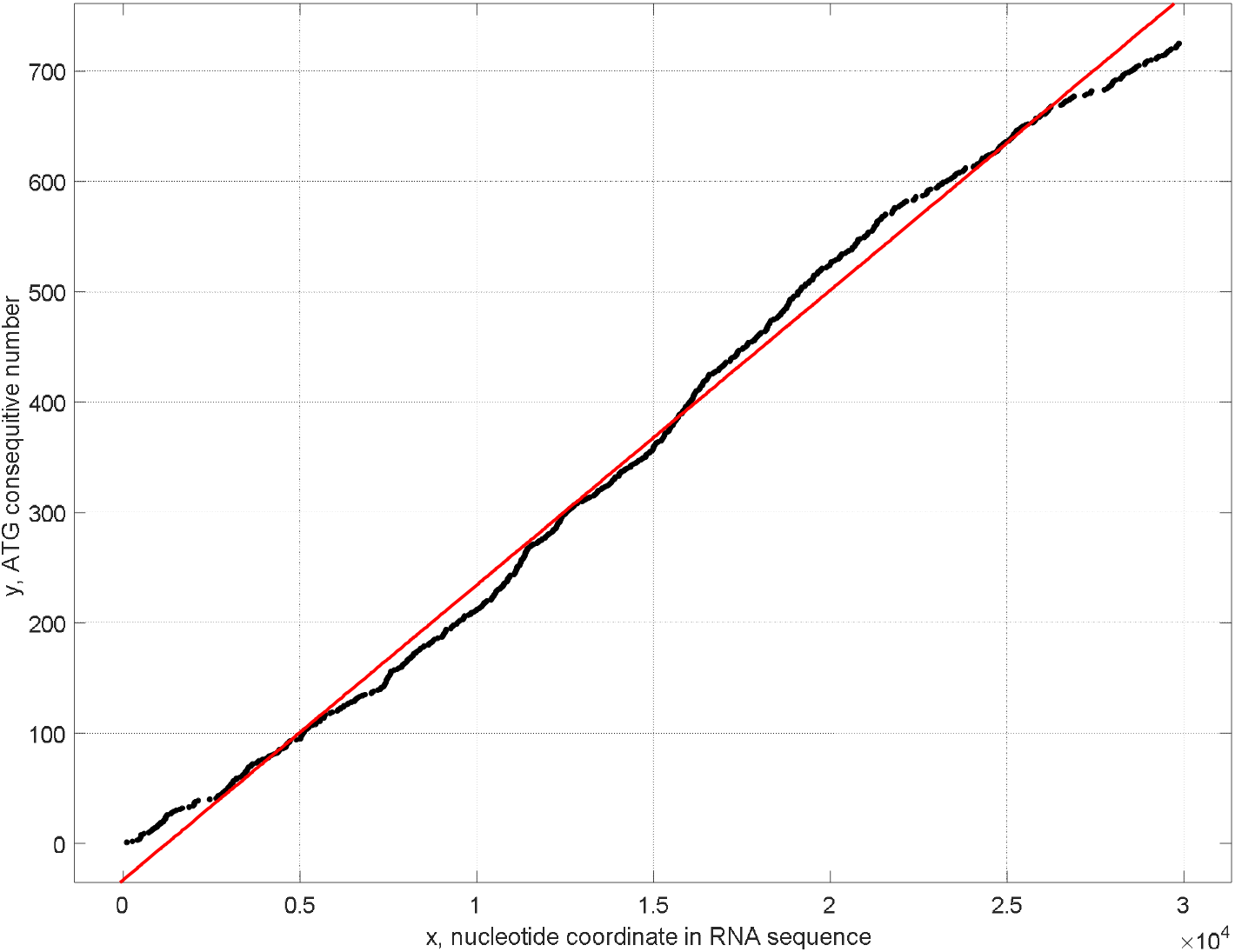
ATG distribution of a complete RNA of SARS CoV-2 virus MN988668.1 (GenBank) and its linear interpolation by formula (1.2).

### 3.3. Comparisons of Averaged Models of Virus RNAs

Today, hundreds of mutated SARS CoV-2 viruses are found every day in different countries. Greek alphabet symbols also designate the most known mutations to GenBank and GISAID data banks [5,6].

Some comparisons of ATG-distributions have already been given in [3]. Here, we provide updated data for newly registered virus mutants to track any mathematically extreme RNA variations. As in [3], we compare the ATG distributions of these mutants with the RNA of the studied above coronavirus MN988668.1 (GenBank). The results are in Fig. 4, where five ATG curves are placed together. An inlet in Fig. 4 shows these curves at the end of these complete genomic sequences on a smaller scale.

**Fig. 4.**
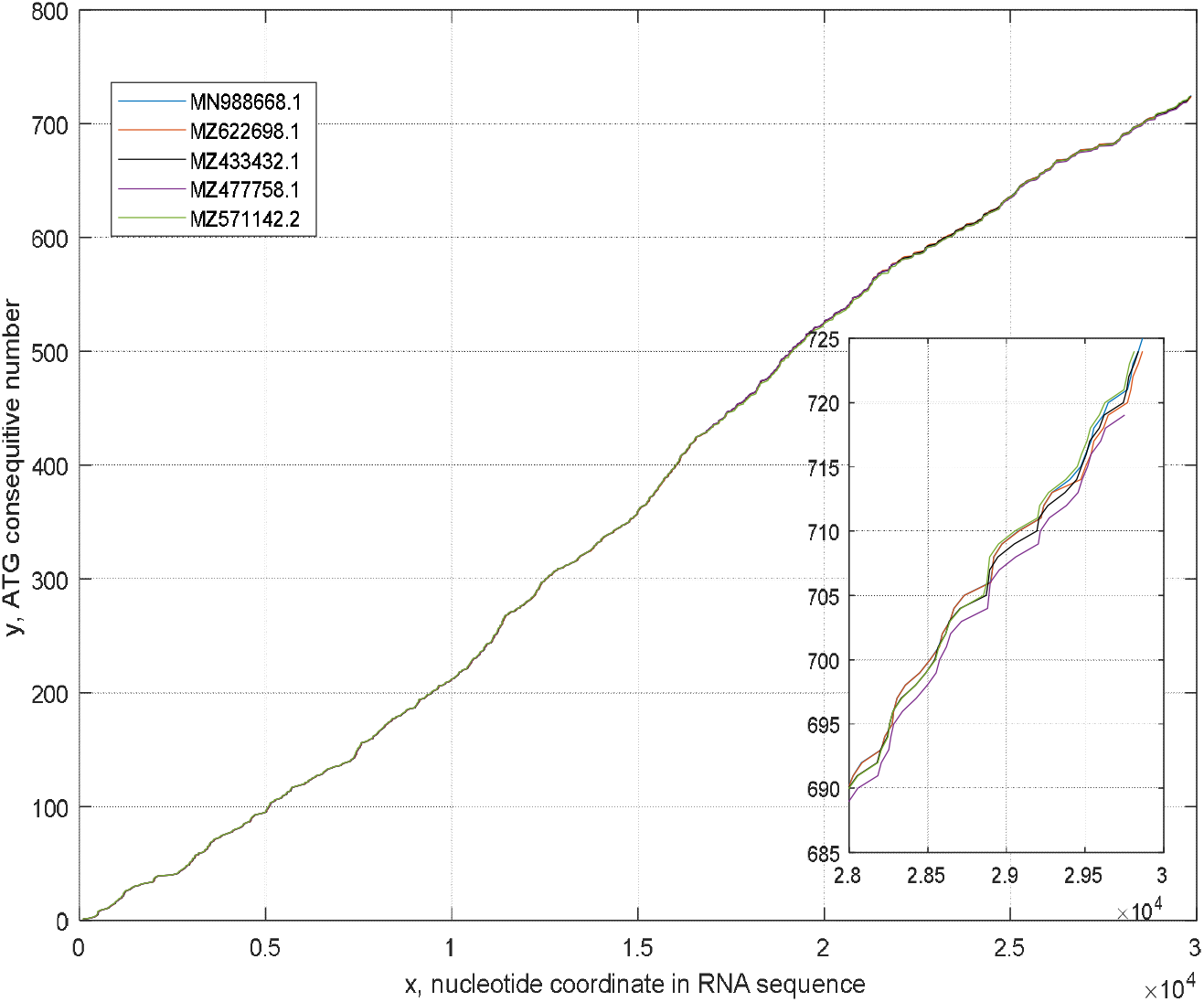
ATG distributions of five RNAs: MN988668.1, MZ622698.1 (‘alfa’, B.1.1.7, registered on 26.07.21), MZ433432.1 (‘beta’, B.1.351, registered on23.06.21), MZ477758.1 (‘gamma’, P.1, registered on 20.07.21), and MZ571142.2 (‘delta’, B.1.167.1, registered on 16.07.21). All are from GenBank [5] where detailed information on these RNAs is placed.

Comparing these ATG curves, it should be noticed that these RNAs have different numbers of non-coding nucleotides at the beginning, and these curves are shifted from each other along the *x* – axis due to that.

Qualitative estimates on the proximity of ATG curves is supported by linear interpolation models, which are

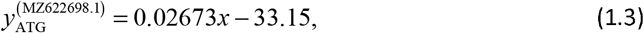

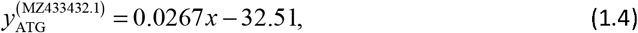

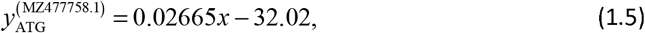

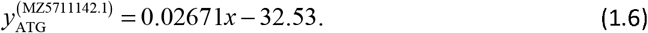

Comparing (1.2) and (1.3)–(1.6), we see relatively strong proximity of these models to each other. It follows that the ATG distributions are prone to be varied, and they can be used as relatively stable schemes of RNAs [3].

### 3.4. Point-to-point Comparison of ATG Distributions

At the same time, we need to notice the limited applicability of averaged models to genomic analysis. Most mutations of slow-rate viruses, to which belong SARS and MERS ones, are with the contents of coding words. Then, perturbations of ATG schemes and distributions of single nucleotides should be studied and coupled with the properties of viruses (see Fig. 2).

We developed a simple algorithm on a detailed comparing of ATG distributions of different virus strains. Each numbered ATG triplet has its coordinate along a sequence. Thanks to some mutations, the length of some coding words is varied together with the coordinate of this triplet. Comparing these varied coordinates with the coordinate of a reference sequence allows estimating some mutations. Researching sequences, we suppose that the error of nucleotide sequencing is negligibly small; otherwise, the results of comparisons would be highly noised.

There are different techniques for comparing sequences known from data analytics [7], including, for instance, calculation of correlation coefficients of unstructured data sequences, comparisons calculating the data distances, clustering of data, and visual analysis [8].

In our case, we calculate the absolute difference between the coordinates of ATGs of the same numbers in the compared sequences. This operation is fulfilled only for the sequences of the same number of ATG triplets; otherwise, excessive coding words are neglected in comparisons. It means, if a compared sequence has several ATG triplets less than the number of ATG one in a reference sequence, the ATGs of reference RNA are excluded from comparisons. Of course, such a technique on comparison geometrical data has its disadvantages, but it allows, anyway, to obtain much information on mutations of viruses in a straightforward and resultative way that will be seen below.

Our approach supposes choosing a reference nucleotide sequence to compare genomic virus data of other samples. Here, it is a complete genomic sequence MN988668.1 from GenBank submitted from Wuhan, China (2020) and studied well in [3]. It has been researched about twenty virus samples from GenBank and GISAID, and some arbitrarily chosen results of comparisons are shown in Figs. 5 and 6.

**Fig. 5.**
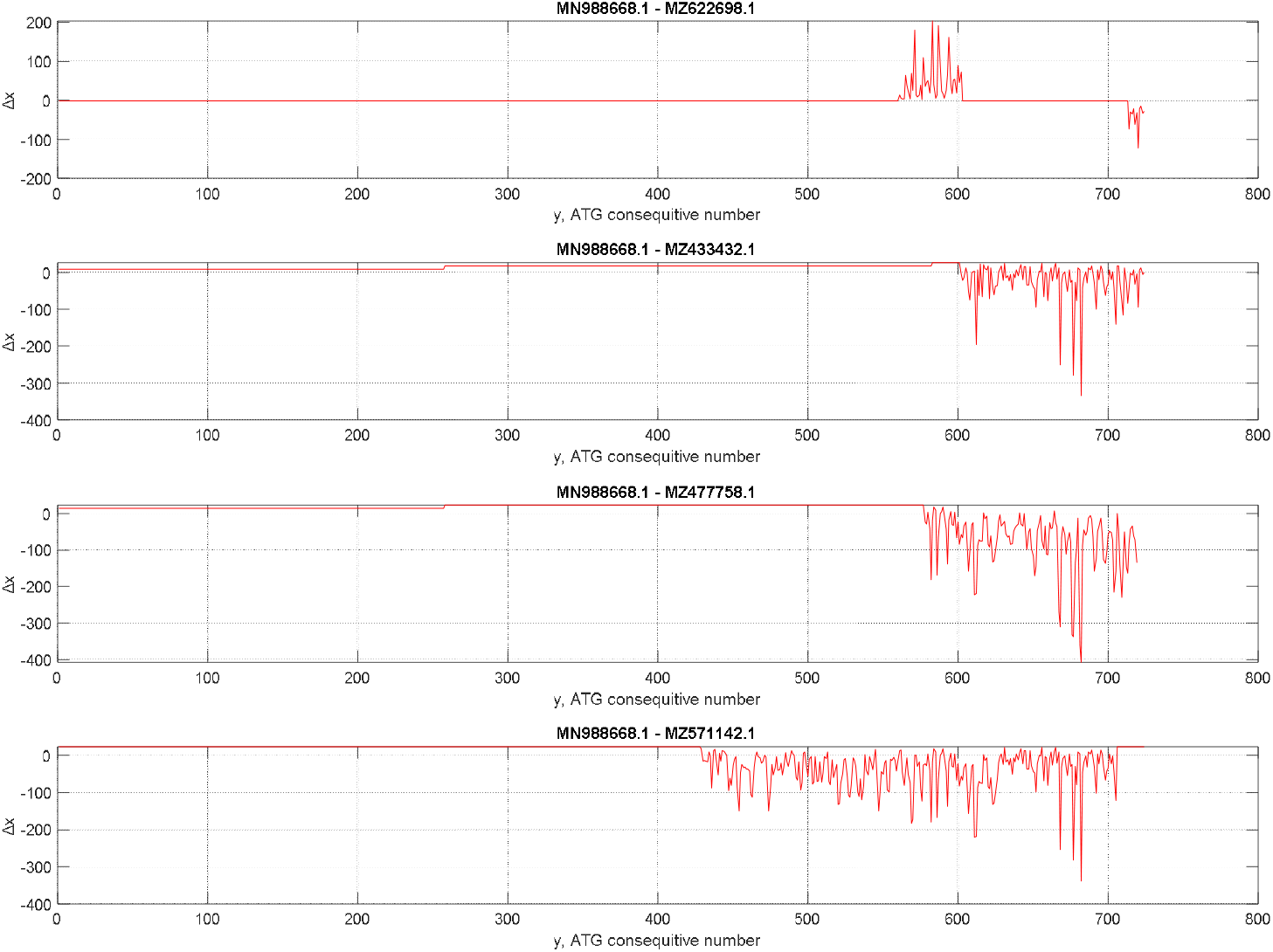
Comparison curves for MZ622698.1, MZ433432.1, MZ477758.1, and MZ571142.1 complete genome sequences of SARS CoV-2 virus samples regarding to the reference sequence MN988668.1. All sequences are taken from GenBank

**Fig. 5.**
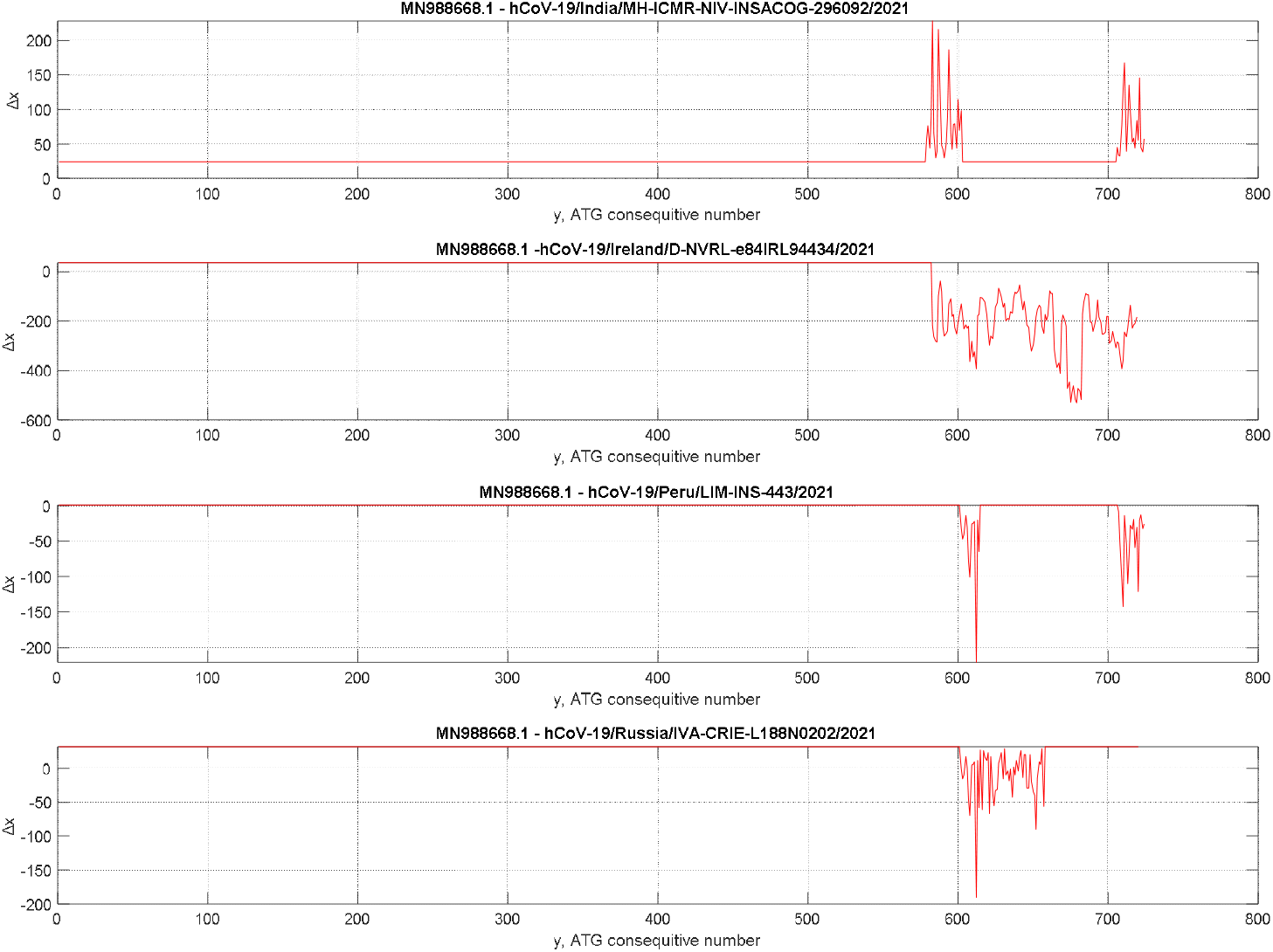
Comparison curves for hCoV-19/India/MH-ICMR-NIV-INSACOG-296092/2021, hCoV-19/Ireland/D-NVRL-e84IRL94434/2021, hCoV-19/Peru/LIM-INS-443/2021, and hCoV-19/Russia/IVA-CRIE-L188N0202/2021 complete genome sequences of SARS CoV-2 virus samples(GISAID [6]) regarding to the reference sequence MN988668.1 (GenBank [5]).

The ordinate axis Δ*x* in these plots shows the deviation of coordinates *x_i_* of ATG triplets from the ATG coordinates of the reference sequence of the same consecutive ATG number *y_i_*. As a rule, due to the different number of non-coding nucleotides at the beginning of sequences, the curves in Figs. 5 and 6 have constant biasing along the Δ*x* axis.

Straight parts of these curves mean that the ATG positions of a compared sequence are not perturbed regarding the corresponding coordinates in the reference RNA, i.e., mutations are only with the variation of coding words without affecting their lengths if these mutations have a place.

In all studied samples, a part of which is shown in Figs. 5 and 6, most perturbations are in the vicinity of the end of *orf1ab* gene, as it is seen using a graphical tool of GenBank [5]. It is detected calculating *x_i_* coordinates according to known *y_i_* and Fig. 4.

It supposed that ATG perturbations could occur in any part of RNA considering the random nature of mutations. Still, to this moment, in the studied samples, only the perturbations have been found showing some steps in the middle of *orf1ab* gene (Fig. 5, curves for the compared MZ433432.1 and MZ477758.1 genome samples).

Interesting to notice that some perturbations of geometry can be self-compensated, and ATG distributions recover their similarity to the reference genome (Fig. 5, a curve for compared MZ622698.1; Fig. 6, curves for the compared hCoV-19/India/MH-ICMR-NIV-INSACOG-296092/2021, hCoV-19/Peru/LIM-INS-443/2021, and hCoV-19/Russia/IVA-CRIE-L188N0202/2021).

Studying geometrical perturbations of ATG distributions and their comparisons with the reference genome, it has been found that these trajectories are individual for the studied samples although it is available mutations without affecting these distributions, and, in these hypothetical cases, the individuality would be lost.

From the studied cases, it follows repeating motives of comparison curves (Figs. 5 and 6). The nature of this is unknown to the authors, but it is not coupled with the lineages of viruses and their clades.

## 4. Conclusion

A new RNA mapping algorithm is applied to build nucleotide and ATG-triplet maps. This big-data code is in calculating Hamming distance of RNA binary symbols and fragments under the search. It allows to get numerical values for the mentioned distance and apply a simple searching algorithm to find the position of each RNA symbol in the complete genomic sequences.

A multi-scale map is built as a set of points of numbered nucleotide positions along a sequence in a grid of lines coordinated with the first symbols of ATG triplets that allow tracking each codon’s content in sequences.

The proximity of ATG distributions of SARS CoV-2 virus samples has been studied in two ways: visual comparisons of ATG curves and comparing coefficients in interpolating polynomials for these curves. This study shows the SARS Cov-2 virus’s slow mutation rate compared to samples obtained in 2020 and 2021.

A specific class of mutations is associated with the change of the coding word lengths. To detect these mutations, the coordinates of ATG triplets have been compared with the coordinates of a reference sequence ATGs. It has been noticed that, to a large extent, the studied RNAs (See Table 1, rows 2-9) have similar distributions with the reference RNA MN988668.1 registered in Wuhan, China. The found most variations of the ATG coordinates are in the vicinity of *orf1ab* gene end.

**Table 1.**
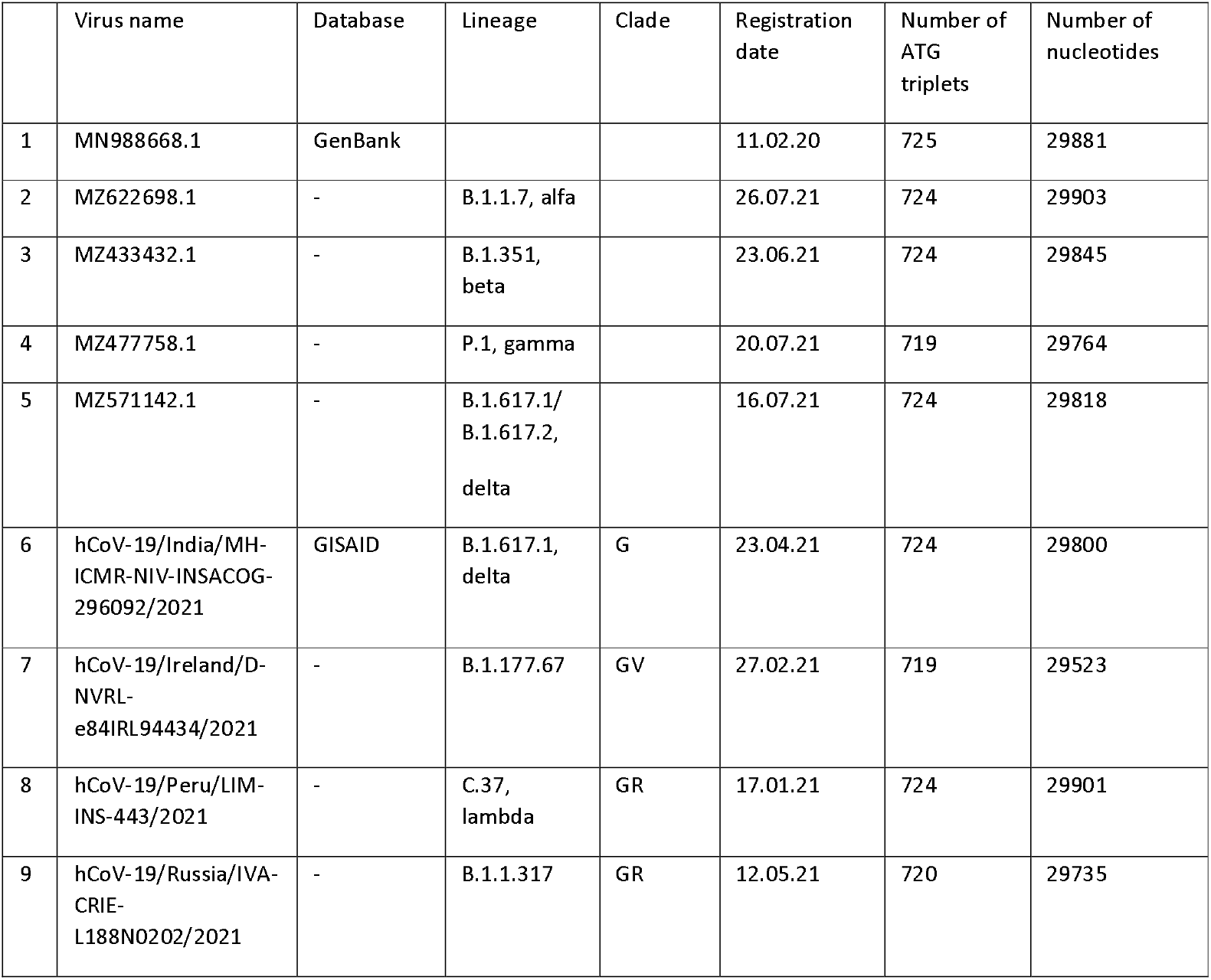
Complete genome sequence data used in this contribution.

Thus, the developed approach on the multi-scale study of virus RNAs has shown effectiveness in analyzing RNA sequences and discovering unknown earlier properties of ATG distributions considered as skeletons of RNAs.

## Abbreviations

SARS CoV-2: severe acute respiratory syndrome coronavirus 2
RNA: ribonucleic acid
DNA: deoxyribonucleic acid.

## Acknowledgments

The authors thank all researchers who placed genomic sequences of viruses in GenBank and GISAID databases.

